# Functional convergence underground? The scale-dependency of community assembly processes in European cave spiders

**DOI:** 10.1101/2023.03.17.533085

**Authors:** Stefano Mammola, Caio Graco-Roza, Francesco Ballarin, Thomas Hesselberg, Marco Isaia, Enrico Lunghi, Samuel Mouron, Martina Pavlek, Marco Tolve, Pedro Cardoso

## Abstract

Understanding how species assemble into communities is a central tenet in ecology. One of the most elusive questions is the relative contribution of environmental filtering versus limiting similarity. Important advances in this area have been achieved by looking at communities through a functional lens (i.e., the traits they express), so as to derive principles valid across species pools. Yet, even using traits *in lieu* of taxonomy, the issue remains controversial because i) environmental filtering and limiting similarity often act simultaneously in shaping communities; and ii) their effect is scale-dependent. We exploited the experimental arena offered by caves, island-like natural laboratories characterized by largely constant environmental gradients and a limited diversity of species and interactions. Leveraging uniquely available data on distribution and traits for European cave spiders, we tested explicit hypotheses about variations in community assembly rules across ecological gradients and scales. We demonstrate that environmental filtering and limiting similarity shape cave communities acting on trait evolution in opposing directions. These effects are strongly scale dependent, varying along multiple environmental gradients. Conversely, the effect of geography on trait composition is weak, indicating that trait turnover in space happens primarily by substitution of species pursuing similar functions due to strong environmental filters. Our findings reconcile contrasted views about the relative importance of the two main mechanisms shaping patterns of biodiversity, and provide a conceptual foundation to account for scaling effects in the study of community assembly.

## INTRODUCTION

An omnipresent scheme in introductory textbooks of ecology illustrates the numerous filters selecting which species end up assembling into local communities from a regional pool. An elusive problem concerning this ‘filtering’ metaphor is quantifying the relative contribution of abiotic and biotic factors in shaping communities (1–3). In a nutshell, environmental filtering is the process whereby abiotic constraints prevent species from establishing in a community, selecting for a narrow set of traits suitable to cope with the local conditions, leading to lower differences in trait composition than expected by chance (“trait underdispersion”). Conversely, biotic interactions such as competition drive functionally similar species to diverge in key phenotypic traits to reduce niche overlap through limiting similarity, leading to higher differences in trait composition than expected by chance (“trait overdispersion”). It follows that looking at biological communities through the lens of functional ecology (i.e., the traits expressed in each community) is one of the most effective ways we have to quantify the interplay between these two assembly processes (4). The use of traits in lieu of species identities allows an explicit focus on the mechanisms generating biodiversity patterns, facilitating the conceptualization of general principles that are valid across species pools.

Even with trait-based approaches, however, it remains difficult to separate the main mechanisms filtering the species pool of potential resident species to the subset that occurs within a given community (α-diversity) and in driving variations across communities (β- diversity) (5). The distinction between environmental filtering and limiting similarity is too often conceptualized as a “black or white” dichotomy, whereby communities are described to be dominated by one or the other process. The ecological reality is instead more nuanced, with the two processes acting simultaneously in shaping communities, although with different intensities given the local conditions (6, 7). Furthermore, like any dimension of biodiversity, functional diversity change is scale-dependent (8, 9), forcing us to account for the pervasive effect that scale has on emerging patterns (10). Since biotic interactions require spatial proximity, the effect of limiting similarity should often decrease with increasing scale and, vice versa, the filtering effect posed by the abiotic environment should increase with spatial scale—generally resulting in a predominance of trait overdispersion at local scales and trait underdispersion at broader scales (11, 12).

Mounting evidence demonstrates how the relative influence of environmental filtering and limiting similarity broadly changes along spatial and temporal gradients—e.g., for vertebrates (8, 11, 13–15) and plants (1, 12, 16). However, there is still controversy on the direction of these changes and their causes (2, 6, 7). To minimize confounding factors and achieve a better understanding of community assembly rules, scientists are therefore increasingly turning their attention to island-like systems (e.g., oceanic islands, lakes, and mountain summits; (17)) and specific biological communities within them [e.g., plants (18, 19); birds (20–22)], as models. The use of island-like systems, i.e., mostly closed, with known histories, and with a relatively low richness of species and interactions, allows ecologists to more easily disentangle community assembly processes while controlling for immigration, extinction, and dispersal dynamics (17, 23, 24).

Under this framework, caves and other subterranean ecosystems stand out as ideal model systems for the study of community assembly processes through a functional lens. Foremost, caves are semi-closed systems extensively replicated across the Earth (25), where stringent environmental conditions promote trait convergence among successful colonizers (26, 27). Second, subterranean communities generally exhibit lower species richness and functional diversity than neighboring surface communities (26, 28, 29) [but see ref. (30)], making it easier to disentangle the relative effect of abiotic conditions and biotic interactions in selecting species possessing specialized traits within the community (31). Third, caves have clear environmental gradients from the surface toward the subsurface (32–34) and display a reduced variability in their abiotic conditions (35), two factors that avoid many of the confounding factors typical of other systems (24).

To study community assembly rules, we leveraged the unprecedented amount of data available for subterranean spiders in Europe (36), namely community composition data for selected caves across the continent (37), and standardized traits for all species (38). A previous analysis of the taxonomic component of this dataset demonstrated a quick turnover in the taxonomic diversity of subterranean spiders across Europe, mediated primarily by geographic distance among caves, and secondarily by the climatic conditions and availability of karst. Conversely, local-scale characteristics of caves exerted a negligible effect on species turnover (39). Here, we explore the functional dimension of these patterns, testing: i) the relative contribution of environmental filtering and limiting similarity in determining community assembly in caves; ii) how functional diversity decays along environmental gradients.

At the α-diversity level, we expect (**H**_**1a**_) communities to be functionally underdispersed because the stringent environmental conditions of caves should filter a narrow set of trait combinations, leading to a less diverse trait composition than what would be expected given taxonomic richness. Concurrently, we predict that (**H**_**1b**_) biotic interactions may exert a significant role where there are fewer available niches (e.g., smaller caves), and when local conditions allow for more contacts among species (i.e., where there is greater habitat connectivity). At the β-diversity level, we hypothesized that (**H**_**2a**_) functional turnover should occur at a lower rate than taxonomic turnover along both geographic and environmental gradients—comparative taxonomic data in ref. (39). This is because we expect that the stringent environmental conditions of caves should act by reducing the volume of the trait space, leading to a functional convergence of cave communities irrespectively of spatial scales. Finally, we predict that (**H**_**2b**_) environmental factors will exert a stronger effect than geographic distance on functional turnover. This is because we expect functional composition to be strongly influenced by local environmental conditions modulating the availability of niches and the potential for interactions.

## RESULTS

### α-diversity

Standard effect size (SES) values for the functional richness of each community were left-skewed, with 63% of caves displaying a prevalence of functional underdispersion over functional overdispersion (Figure 2a, 2b). Still, most of these caves clustered towards SES values close to zero (Figure 2b), with only seven communities completely underdispersed and one completely overdispersed (p < 0.05). In general, caves with a prevalence of overdispersion were concentrated at southern latitudes, especially in the Dinaric karst (western Balkans) (Figure 2a). A generalized least squares model fitted through the data suggested that communities in caves with a greater depth (negative drop), occurring within larger karst areas, and in areas with a broader annual temperature range were more likely to be functionally overdispersed (Figure 2c).

### β-diversity

Patterns of functional β-diversity were primarily driven by a few environmental gradients (Figure 3). Most of the variation in β-diversity was due to replacement of trait space among communities (β_replacement_; Figure S1), with patterns largely mimicking the variation in total β- diversity (β_total_). Conversely, the contribution of β_richness_ was negligible in all cases except for cave drop (Figure S2).

**Table 1.** Estimated regression coefficients for the generalized least square model. Significant values are highlighted in bold. LGM = Last Glacial Maximum.

| TERM | ESTIMATE | STANDARD ERROR | T-STATISTIC | P-VALUE | VIF<sup>1</sup> |
| --- | --- | --- | --- | --- | --- |
| INTERCEPT | −0.23 | 0.05 | −5.02 | **> 0.001** |  |
| ENTRANCE SIZE<br>[m<sup>2</sup>] | −0.01 | 0.05 | −0.16 | 0.870 | 1.17 |
| DEVELOPMENT<br>[m] | −0.02 | 0.05 | −0.38 | 0.707 | 1.27 |
| NEGATIVE DROP<br>[m] | 0.11 | 0.05 | 2.16 | **0.031** | 1.21 |
| ELEVATION<br>[m] | 0.1 | 0.05 | 1.77 | 0.077 | 1.35 |
| LGM ICE DISTANCE<br>[km] | 0 | 0.06 | 0.01 | 0.994 | 1.3 |
| KARST AREA<br>[km<sup>2</sup>] | 0.12 | 0.05 | 2.33 | **0.020** | 1.22 |
| TEMPERATURE<br>[°C] | 0.06 | 0.06 | 0.91 | 0.364 | 1.86 |
| ANNUAL RANGE<br>[°C] | 0.26 | 0.06 | 4.22 | **> 0.001** | 1.66 |
| PRECIPITATION<br>[mm] | 0.01 | 0.06 | 0.14 | 0.889 | 1.7 |

<sup>1</sup> Variance Inflation Factor.

The effect of geographic distance on functional β-diversity followed a power-law curve (linearly asymptotic), and was rather weak—that is to say, at increasing distance between two caves, there was only a limited turnover in functional richness (Figure 3a). Interestingly, when looking at variation in the effect over the geographic gradient (Figure 3c), we observed a prevalence of trait overdispersion at a smaller spatial scale which progressively decreased toward zero when caves were >2000 km apart.

Environmental predictors identified as important by the BBGDM were, in order of importance, cave development, elevation, precipitation, entrance size, and annual range of temperature. The contribution of additional predictors was negligible. The rate of turnover along the cave development gradient was monotonically asymptotic, with rates of turnover steeply increasing in the first portion of the gradient before reaching a plateau (Figure 3a). This effect was significant along the whole gradient (Figure 3c). We also observed some degree of turnover along the gradients of elevation, precipitation, and temperature range. That is, communities in caves with different elevations, temperatures, and precipitation regimes tend to express different functions. For precipitation, SES values indicated that there is underdispersion along the first half of the gradient and an increasing predominance of functional overdispersion in the second half of the gradient. The pattern was reversed for the annual range of temperature.

## DISCUSSION

Focusing on the underexploited natural laboratory offered by caves, and relying on an unprecedented data baseline in the context of subterranean biology, we studied functional diversity patterns in subterranean spider communities across Europe, testing general hypotheses ruling community assembly. Two important points, largely generalizable across systems and species pools, emerge from our analysis.

The first key point is that environmental filtering (causing trait convergence) and limiting similarity (causing trait divergence) are not mutually exclusive processes (40). Even in caves, ecological systems where environmental filtering is meant to be particularly strong (28), the relative influence of these two processes varied substantially given the local habitat conditions. Whereas the direction of SES for functional richness was predominantly towards underdispersion (Figure 1b), the majority of values were close to zero, meaning environmental filtering and limiting similarity were both acting in equally weak or strong, but opposing, directions. Environmental filtering is indeed a demonstrably strong factor in caves, with many traits and parts of the potential functional space being absent. Yet, our results add quantitative evidence to a growing body of literature (27, 30, 41, 42) emphasizing the importance of reconsidering the role of species interactions (especially competition) as an important force driving the evolution of cave communities.

**Figure 1.**
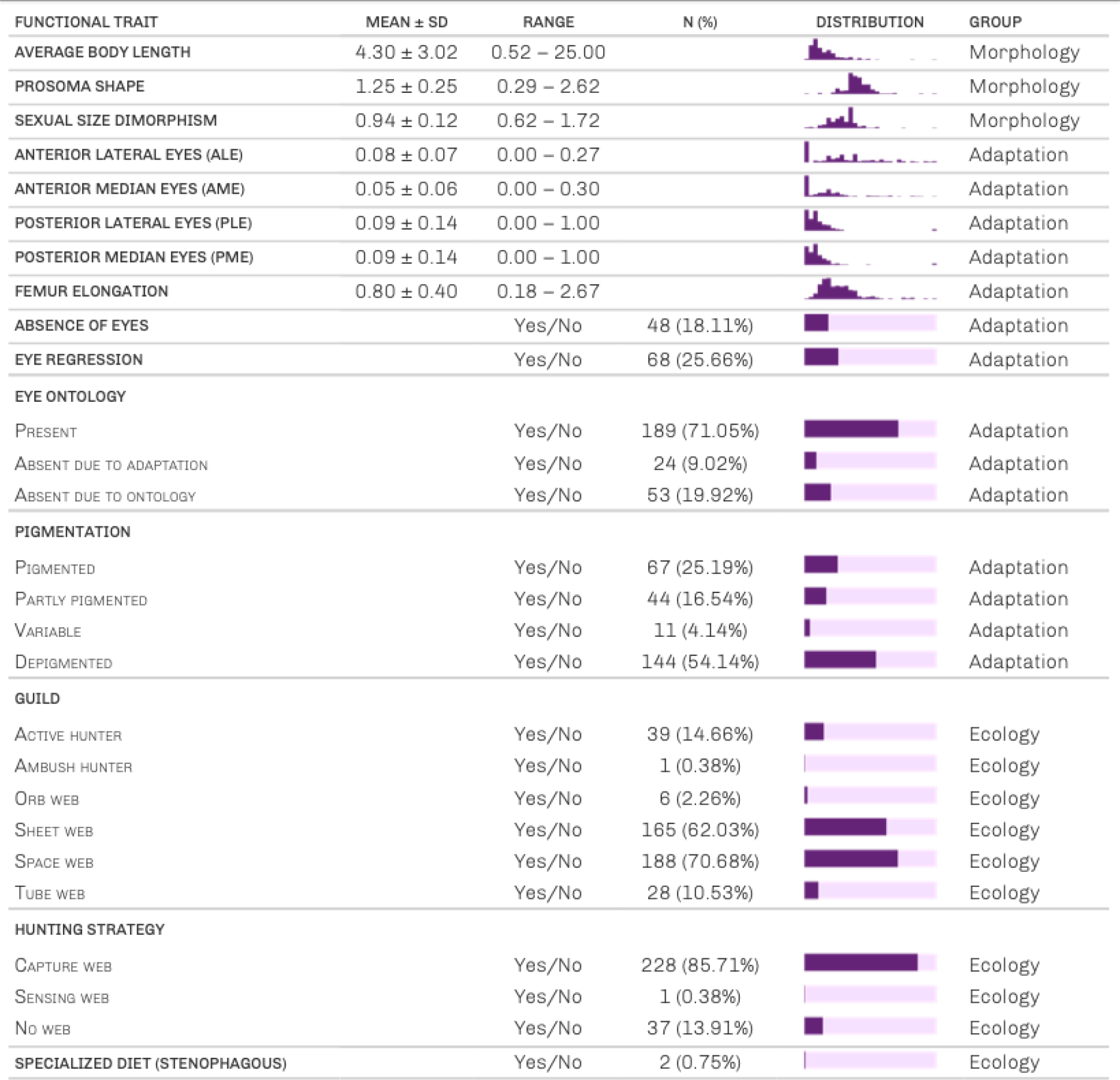
Summary of the traits used in the analysis. We refer to ref. (38) for a full description of traits and their hypothesized functional meaning. Column “Group” refers to the grouping used in the estimation of weights for the Gower distance *sensu* ref. (79), whereby: “Adaptation” are traits related to morphological adaptation to subterranean conditions (especially darkness); “Morphology” are traits describing general morphology of species; and “Ecology” refers to traits describing webs and hunting strategies.

Subterranean communities with local trait overdispersion were more frequently associated with large karst patches, areas with broader temperature ranges, and deeper caves (Figure 2c), all conditions that provide more niche space to be exploited. This was particularly evident in the Dinaric karst (western Balkans), the most important global hotspot of subterranean biodiversity (43, 44), where virtually all cave trait spaces were predominantly overdispersed. Indeed, large patches of karst, such as in the Dinarides, implies greater habitat availability (45) and possibly connectivity (46), hence a higher niche space, and a greater chance of contact among species. At the same time, the positive association between trait overdispersion and temperature range can be interpreted in the light of the influence of temperature variability on species range size and dispersal (47–49). As demonstrated for subterranean spiders in the genus *Troglohyphantes* (50), areas with larger temperature seasonality tend to allow greater overlap among species ranges, hence increasing local diversity and the likelihood for species coexistence. Finally, communities in caves with a greater drop tend to be, on average, predominantly overdispersed. Deeper caves tend to express more areas with differing availability of resources, offering more possibility for different communities with contrasting traits.

**Figure 2.**
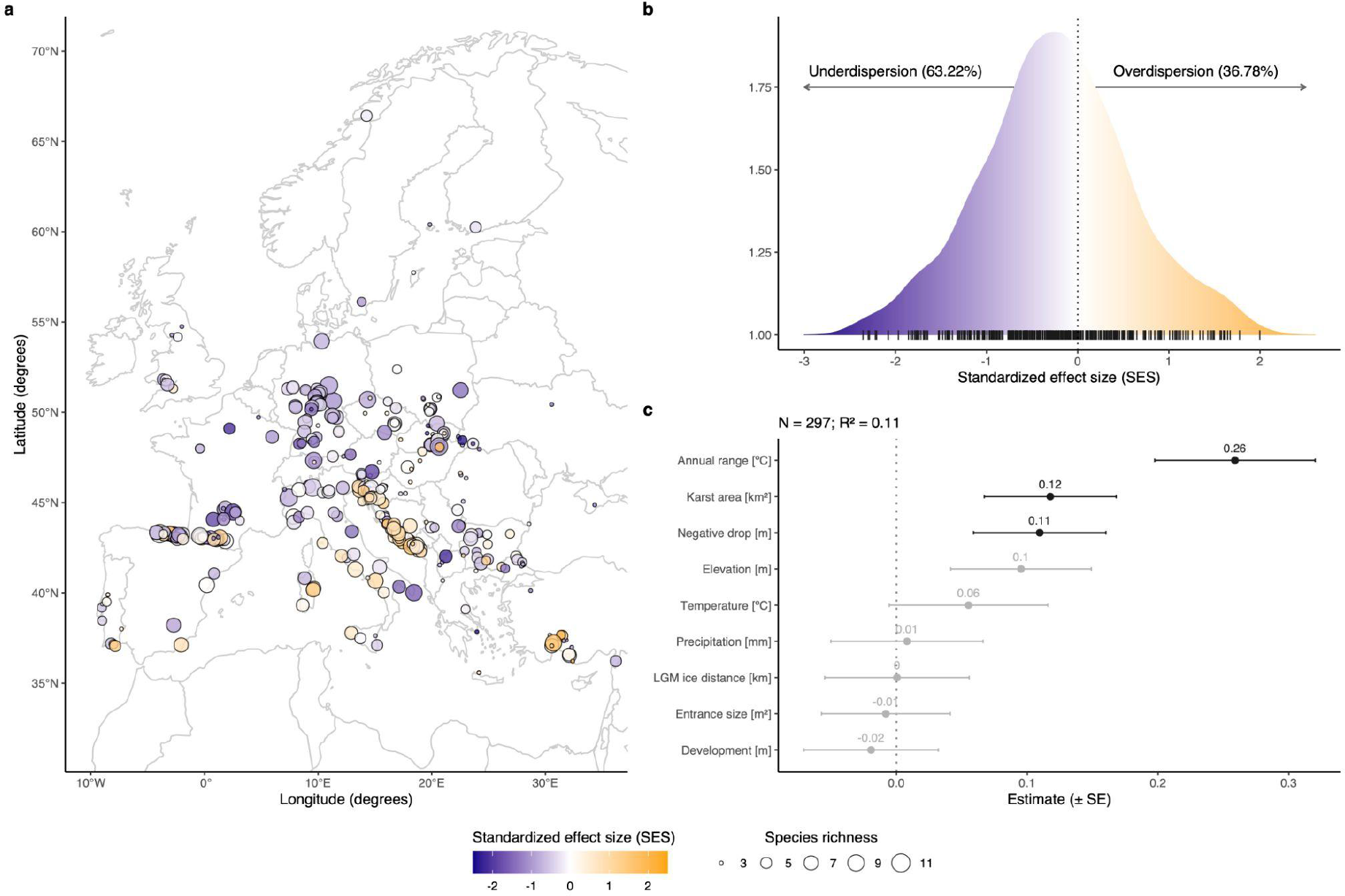
Functional diversity at the α-diversity level for subterranean spider communities in Europe. **a**) Distribution of the studied caves (N = 367 caves, after excluding caves with species richness < 3). The size of each dot represents species richness. Dots are colored according to their standard effect size (SES) value for functional richness, where functional richness is estimated as the volume of the hypervolume representing each cave’s trait space. **b**) Density of SES for functional richness across the studied caves. Percentage of caves with negative or positive SES are indicated. Dark lines at the bottom of the density curve show the frequency of observed values. **c**) Environmental factors driving variation of SES values for functional richness. Estimated parameters are based on a generalized least square model (significant effect in a darker color). Error bars indicate standard errors. The exact estimated regression parameters and p-values for the model are in Table 1. Note that the sample size of this model is 297 (not 367) because of missing data in the environmental data for some caves. LGM = Last Glacial Maximum.

**Figure 3.**
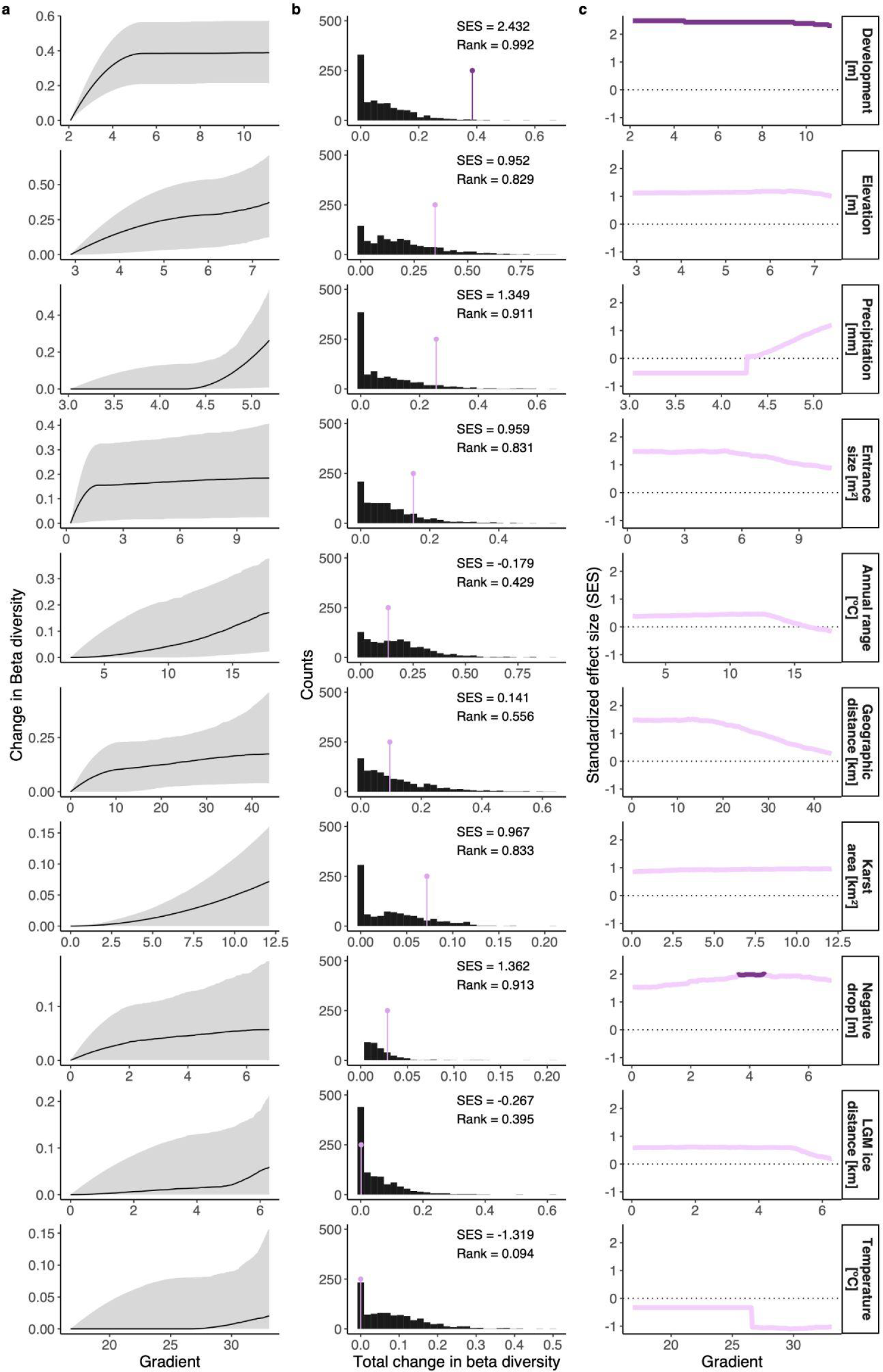
Results of Bayesian bootstrap generalized dissimilarity modeling for change in total functional β-diversity (β_total_) of subterranean spider communities across Europe (i.e., unit increase in mean β along a given gradient). Variables are sorted by their contribution (on top: highest contribution). **a**) Fitted I- splines (partial regression fits) for the considered environmental and geographic gradients. The maximum height reached by each curve indicates the total amount of compositional turnover explained by that variable (holding all other variables constant), whereas the shape of each spline indicates how the rate of compositional turnover varies along the gradient. **b**) Distribution of expected values (histogram) versus the observed value (colored line) of each environmental and geographic gradient, based on null modeling (999 iterations). In other words, these panels provide information as to whether the effect of a given variable is higher or smaller than expected given taxonomic composition. **c**) Variation in the magnitude of the standard effect size (SES) value along the observed gradient. In other words, these panels provide information as to whether the effect of a given variable in determining trait dispersion changes along the gradient. In **b** and **c**, significant effects (Rank < 0.025 | > 0.975) are highlighted with a darker purple.

The second key point emerging from our study is about the importance of scale in the perception of community assembly patterns. Accounts for scale in trait analyses have been achieved, for example, by looking at variations in individual trait values along ecological gradients [e.g., elevation (51)], or by contrasting taxonomic and functional diversity change in highly dispersive organisms—birds (8). Here, we devised a novel approach to account for the magnitude of trait dispersion change along the studied ecological gradients, combining gradients and traits all together in a single model. We observed how functional β-diversity patterns varied along multiple ecological gradients. The most important one was the difference in the development among the studied caves, whereby the largest replacement of functions occurred between pairs of caves with divergent development (i.e., large versus small caves; with an inflection point with cave >10 meters in development). A plausible explanation is that cave development is a proxy for the availability of spatial niches. In particular, small caves will be primarily colonized by spiders adapted to the cave entrance conditions, and large caves will often sustain a greater number of specialized species, accounting, overall, for drastically different functions. Other important gradients of variations were elevation and precipitation, reflecting the influence of climatic conditions and habitat heterogeneity in the local structuring of functions.

In terms of distance decay in functional diversity, replacement of function was not as pronounced as the replacement of species (taxonomic distance decay) observed in ref. (39) for the same set of caves. This means that, although some replacement of traits do occur, overall turnover happens by substitution of species pursuing similar functions. Still, when decoupling functional patterns from taxonomic diversity, the functional responses varied along the geographic gradient according to theoretical expectations, showing stronger overdispersion at smaller distances, and progressive moving toward SES values of zero at larger distances. This highlights the scale dependency of regional trait dispersion, with nearby caves more likely to have interacting communities and such effect becoming weaker with increasing spatial distance.

## CONCLUSIONS

The use of caves as model systems for investigating (macro-)ecological patterns in space and time is still underexploited (52). This is partially a problem related to the objective difficulties of working in caves (resulting in a general lack of data at the right resolution) and partly a methodological problem. Nonetheless, thanks to the recent development in databases of species distributions and traits, and the emergence of novel analytical tools, there is a vast potential to leverage these systems as ideal settings in which to model across space. Using an explicit functional diversity approach, we showed that i) even systems with stringent environmental conditions maintain the potential for trait differentiation via biotic interactions, especially in areas of greater habitat connectivity; and ii) the relative influence of environmental filtering and limiting similarity change with scale, along clear ecological gradients. Overall, our findings reconcile contrasted views about the relative importance of the two main mechanisms shaping patterns of biodiversity, and provide a conceptual foundation to account for scaling effects in the study of community assembly. This information is key amidst escalating global anthropogenic threats (53), insofar as realistic predictions of biodiversity change require explicitly accounting for community assembly processes (54).

## MATERIAL & METHODS

### Community-level data

We obtained data for subterranean spider communities across Europe from ref. (37), which we refer to for a full account of the dataset and the methods used to assemble it. The dataset comprises data from 475 subterranean sites (limestone, volcanic, talus, and salt caves, as well as artificial sites including mines, blockhouses, and cellars; the general term ‘cave’ is used hereafter) across 27 European countries, covering a latitudinal range from 35° to 70°. The dataset only includes subterranean sites for which spider fauna is exhaustively known. For each site, the spider composition is represented as incidence data—presence/absence of each species. Note that we focused solely on “subterranean spiders” (36, 38), excluding “accidental” surface species (55) occasionally found underground.

### Environmental gradients

We collated a site-by-environment matrix including local-scale environmental characteristics of each cave and broad-scale variables extracted from rasters using the coordinates of the cave entrance. A full description of both local- and broad-scale variables is given in refs. (37, 39).

As local-scale predictors, we used the altitude of the cave entrance (in meters a.s.l.), the main entrance size (a numerical estimation of the dimension of the main entrance in square meters), cave development (total planimetric development of the cave in meters), and cave depth (total drop in meters). These are frequently used variables in macroecological analyses focused on caves (56), which we here interpreted as proxies for local-scale conditions and niche space availability. For example, caves with a vertical drop and a large entrance tend to accumulate more external food resources (detritus) than horizontal caves with a very narrow entrance.

As broad-scale predictors, we included three climatic variables (mean annual temperature, annual temperature range, cumulative precipitation), one variable reflecting availability of carbonatic rocks (karst), and one biogeographical factor (the distance of each cave to the margin of the glacier in Last Glacial Maximum; ca. 21,000 years ago). We extracted climatic data from WordClim 2 rasters (57) at a resolution of 2.5 minutes. Although they may fail-short to capture microclimatic variability within caves (58), these broad-scale variables are good surrogates for general subterranean climatic conditions (59–62). We extracted the size of the karst patch in which a cave occurs using the World Map of Carbonate Rock Outcrops (version 3.0). Given that most locations in our database were karst caves, we interpreted this variable as a good proxy of habitat availability in the surrounding of each cave, and an indirect measure of habitat connectivity (45, 46). Finally, we derived the distance of each cave from the Last Glacial Maximum glacier from reconstructions by ref. (63). We interpreted this as a *proxy* for the influence of past glacial cycles on the current distribution of subterranean species (64, 65).

### Functional traits

For each spider species included in the database, we derived functional traits from ref. (38), which we refer to for a full description of the trait data matrix and data collection methods. For the purposes of the analysis, we selected a subset of traits from the whole trait matrix, representing: i) general morphology of species; ii) morphological adaptation to subterranean conditions; and iii) webs and hunting strategies (Figure 1). To ensure exact matching between the spider species names in the community and trait matrices, we standardized and updated taxonomy using the function *checknames* in the R package ‘arakno’ version 1.1.1. (66).

### Data analysis

We analyzed data in R version 4.1.2 (67), using the suite ‘tidyverse’ (68) for data manipulation and visualization. In all functional diversity analyses, we followed the general analytical pipeline described in ref. (69), and the protocol for transparent reporting by ref. (70). A reproducibility checklist for the study is available in Table S1. Since functional analyses were computationally demanding, we ran all analyses in high-performance computing services (see “**Acknowledgments**”).

#### Data exploration

We carried out data exploration following refs. (70, 71), checking variable distribution, multicollinearity, and the presence of missing data (Figure 1). As a result of data exploration, we standardized all continuous traits (mean = 0 and standard deviation = 1) to ensure comparable ranges among different traits. In the environmental matrix, we checked variable distributions and log-transformed all numerical variables (except coordinates, annual temperature range, and mean temperature) to homogenize distribution and reduce the effect of outliers. None of the predictors showed correlation values higher than Pearson’s *r* > ±0.7 (71).

#### Functional space estimation

We estimated the trait space of each cave using probabilistic hypervolumes (72–74). Probabilistic hypervolumes have two key advantages over other commonly used trait-space characterizations [e.g., dendrograms (75) or convex hulls (76)]: i) they allow the detection of areas of higher or lower density in the trait space, thus representing uneven probabilities of finding a species with a given trait combination throughout the boundaries of the trait space; and ii) they are less sensitive to outliers than convex hulls (74, 77).

Prior to analyses, we filtered caves with less than three species because these might lead to uninformative trait spaces, resulting in a total sample size of 367 caves. Since the trait matrix was a mixture of continuous, binary, and categorical traits, and contained some missing data for certain traits, we used a Gower distance to estimate trait dissimilarity among species (78). Because different traits were broadly associated with different functional meanings, we used the optimization method by ref. (79) to attribute weight to traits within the three groups of variables (column “grouping” in Figure 1). We analyzed the resulting distance matrix through Principal Coordinate Analysis with the R package ‘ape’ version 5.5.0 (80), extracting three orthogonal axes that we used to delineate the probabilistic hypervolumes for each cave. Using three trait axes ensures a good trade-off between accuracy and computation time (9, 81). We constructed hypervolumes with a Gaussian kernel density estimator and a default bandwidth for each axis (82), as implemented in the function *hypervolume_gaussian* in the package ‘hypervolume’ version 3.0.1 (83).

#### Calculation of α- and β-diversity

We measured the properties of the estimated trait spaces using hypervolume-based functions (74) from the R package ‘BAT’ version 2.7.1 (84, 85). We calculated the functional richness of each community (α-diversity) as the total volume of each hypervolume (*kernel*.*alpha* function). We estimated pairwise functional β-diversity among communities as a Sørensen dissimilarity index, calculated through a modified version of the *kernel*.*beta* function that enables parallel estimation of pairwise comparisons (9). This estimation of β-diversity further decomposes the two processes underlying overall dissimilarity (β_total_) among hypervolumes following ref. (86), namely: the replacement of trait space between communities (β_replacement_), and the net differences between the amount of trait space enclosed by the two communities (β_richness_). β-diversity ranges from 0 (identical trait spaces) to 1 (fully dissimilar trait spaces).

#### Null modeling

Estimations of functional diversity are dependent on the taxonomic structure of the communities. Statistically controlling for this association may reveal the actual degree of importance of trait composition to community patterns (69, 87). To this end, we randomized the rows of the trait matrix 999 times to generate a null distribution of each hypervolume-based trait space. For each random iteration, we calculated all α- and β-diversity measures (see the next section). We expressed the deviation of observed values from the null distribution as standard effect sizes (SES). Because standardized effect sizes may lead to biased conclusions if null values show an asymmetric distribution or deviate from normality, we estimated non-parametric effect sizes using probit transformed p-values (12). Probit transformation is used as an alternative to logit transformation in generalized linear models to transform probabilities into the minus-infinity-to-infinity range (88). This approach is known to partially underestimate the effect size when the observed value is completely outside the null distribution; however, this problem was trivial in our case, as none of our observed values fell outside the null distribution (that is, p-value = 0 or 1).

#### Hypothesis testing

To test our first set of hypotheses on alpha diversity patterns (**H**_**1**_), we modeled the relationship between SES values for functional richness (α-diversity), and all local and broad-scale environmental characteristics of each cave using a generalized least squares fitted with the package “nlme” version 3.1-157 (89). To account for spatial autocorrelation, we introduced an exponential correlation structure on the longitude and latitude coordinates of each cave. Prior to model fitting, we standardized all predictors (mean = 0 and standard deviation = 1) to ensure comparability among effect sizes. We validated the model by inspecting the normality of residuals, heteroskedasticity, and degree of collinearity (71).

To test our second set of hypotheses on beta diversity pattern (**H**_**2**_), we used a Bayesian bootstrap extension of generalized dissimilarity modeling (BBGDM), as implemented in the R package ‘bbgdm’ version 1.0.1 (90). Generalized dissimilarity modeling is a matrix regression technique that incorporates variation in the rate of compositional turnover along an environmental or spatial gradient (non-stationarity) in a monotonic nonlinear fashion (91, 92). Because the elements of a dissimilarity matrix are not fully independent, BBGDM uses a Bayesian bootstrap procedure to correct the uncertainty of model parameters (90). We used as input the predictors and the functional β-diversity matrices. We fitted individual BBGDMs for the three functional β-diversity matrices (β_total_, β_replacement_, and β_richness_) with default parameters of three I-splines for each predictor and default knot values.

Because we ran BBGDM for both the actual β-diversity matrices and also the β- diversity matrices resulting from null trait matrices (see section “*Null modeling”*), several metrics of SES could be derived to address different questions. Here, we tested whether a given variable had a stronger or weaker effect than what would be expected given taxonomic richness by extracting the sum of splines coefficients for each variable in the 999 BBGDMs. This way, we generated a null distribution of model coefficients which could further be tested using non-parametric SES. The sum of spline coefficients of each variable describes the total change in β- diversity promoted by a single predictor holding all other predictors constant. Furthermore, we tested whether the SES of a predictor changes along the environmental or geographic gradient. This is relevant to evaluate if communities show a stronger signal of overdispersion (positive SES) when caves are in close geographic proximity, and a stronger signal of overdispersion (negative SES) when caves are far apart. To this end, we extracted the prediction values for all the sites in each of the null BBGDMs and used these to generate a null distribution of prediction values which was further compared against the observed predictions and converted into SES values.

## SUPPLEMENTARY MATERIALS

Functional diversity protocol checklist

Figure S1

Figure S2

Supporting literature

## DATA AVAILABILITY

Data used in the study are available in Figshare (Community data: https://doi.org/10.6084/m9.figshare.8224025.v1; Trait data: https://doi.org/10.6084/m9.figshare.16574255). Metadata and data collection methods are described at length in ref. (37, 38). Climatic rasters are available from WordClim 2 (57), last glacial maximum reconstruction from ref. (63), and carbonate rocks maps from the World Map of Carbonate Rock Outcrops (version 3.0). A cleaned version of the rasters is also available on GitHub (see Code Availability), including the code for data cleaning.

## ACKNOWLEDGEMENTS

This project has received funding from the European Union’s Horizon 2020 research and innovation program under the Marie Sklodowska-Curie grant agreement No 882221 and the Biodiversa+ project DarCo. CG-R was funded by Finnish Cultural Foundation. The authors acknowledge the support of NBFC to CNR, funded by the Italian Ministry of University and Research, P.N.R.R., Missione 4 Componente 2, “Dalla ricerca all’impresa”, Investimento 1.4, Project CN00000033. The authors wish to acknowledge CSC – IT Center for Science, Finland, for computational resources. Finally, a special thanks goes to Federico Riva for critical feedback on the manuscript.

## AUTHOR CONTRIBUTIONS

SMa, CG-R, and PC conceived the idea. SMa and CG-R designed methodology. All authors except SMa, PC, and CG-R collected trait data. CG-R led the analysis, with support by SMa. SMa wrote the first draft, with support by CG-R. All authors contributed to the writing with suggestions and critical comments.

## COMPETING INTERESTS

None declared.

## Notes

https://doi.org/10.6084/m9.figshare.8224025.v1

https://doi.org/10.6084/m9.figshare.16574255

### Competing Interest Statement

The authors have declared no competing interest.

### Summary of Updates

Wrong affiliation for one author

