## Appendix S1 for "Functional convergence underground? The scale-dependency of community assembly processes in European cave spiders"

**TABLE OF CONTENT**

Table S1

Figure S1

Figure S2

Supporting literature

**Table S1.** Functional diversity protocol checklist *sensu* ref. (1).

| **Field** | **Response** |
|---|---|
| Study title | Functional convergence underground? The scale-dependency of community assembly processes in European cave spiders |
| Authors | Stefano Mammola, Caio Graco-Roza, Francesco Ballarin, Thomas Hesselberg, Marco Isaia, Enrico Lunghi, Samuel Mouron, Martina Pavlek, Marco Tolve, Pedro Cardoso |
| Hypothesis | We tested: i) the relative contribution of environmental filtering and limiting similarity in determining community assembly in caves; ii) how functional diversity decays along environmental gradients. |
| Ecological unit | Subterranean systems (caves) |
| Power analysis | No |
| Focal taxa | Spiders (Arachnida: Araneae) |
| Resolution | Species level |
| Number of taxa | 326 |
| Sampling unit | Sites (subterranean habitats) |
| Number of sampling units | 475 |
| Sampling effort | 475 |
| Occurrence data type | Presence-only |
| Number of traits | 20 |
| Continuous traits used | 8 |
| Discrete traits used | 0 |
| Binary traits used | 9 |
| Fuzzy-coded traits used | 3 |
| Trait resolution | Coarse |
| Sample size per species and trait | Species level ; Mean value at the species level |
| Hypothesized function of | Traits related to subterranean adaptation, ecology, and hunting |

| | |
|---|---|
| each trait | strategies |
| Intraspecific variation accounted for? | No |
| Data source | Online database |
| Data exploration | Data visualization, Collinearity assessment, Missing data assessment, Species sampling coverage |
| Collinearity assessed? | Several traits related to subterranean adaptation. We used gower distance and principal coordinate analysis to extract three trait axes for analyses. |
| Transformations done? | Scaled and centered continuous traits |
| Missing data accounted for? | Yes |
| Imperfect detection control | No |
| Functional trait space method | Probabilistic hypervolume |
| Dissimilarity metric used for trait space (if applicable) | Gower |
| Level of analysis | Alpha diversity (within group), Beta diversity (between group) |
| FD method | Richness |
| Model | GLS (for alpha) ; BBGDM (for beta) |
| Effect sizes | Reported in Figure 2 and 3. |
| Model support | $R^2$ |
| Model uncertainty | Confidence interval |
| Validation method | Normality of residuals, heteroskedasticity, degree of collinearity (VIF) |
| Preregistration | No |
| Code link | https://github.com/StefanoMammola/ Mammola_et_al_Macroecology_spider_traits.git |
| Community data link | https://doi.org/10.6084/m9.figshare.8224025.v1 |
| Trait data link | https://doi.org/10.6084/m9.figshare.16574255 |
| Environmental data link | World Map of Carbonate Rock Outcrops; Wordclim 2; clean raster |

| | |
|---|---|
| | also available in GitHub: https://github.com/StefanoMammola/Mammola_et_al_Macroecology_spider_traits.git |
| Data and Code description | README file; software and package version numbers; commented code. The code we provided can reproduce all results, figures and tables. |
| Date | 16 June 2022 |

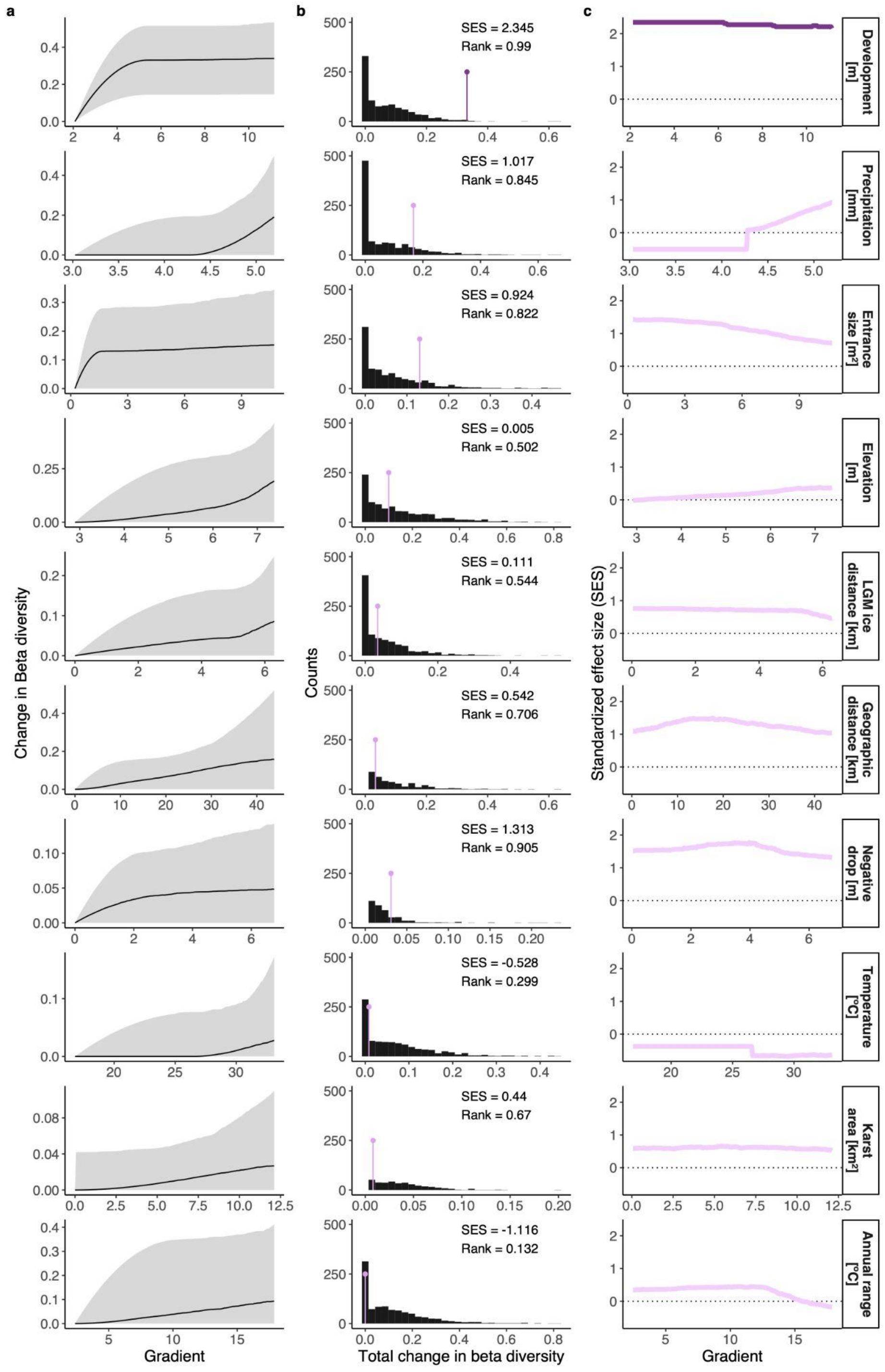


**Figure S1.** Results of Bayesian bootstrap generalized dissimilarity modeling for change in the replacement component of functional β-diversity ($\beta_{replacement}$) of subterranean spider communities across Europe. Variables are sorted by their contribution (on top: highest contribution). **a**) Fitted I-splines (partial regression fits) for the considered environmental and geographic gradients. The maximum height reached by each curve indicates the total amount of compositional turnover explained by that variable (holding all other variables constant), whereas the shape of each spline indicates how the rate of compositional turnover varies along the gradient. **b**) Distribution of expected values (histogram) versus the observed value (colored line) of each environmental and geographic gradient, based on null modeling (999 iterations). In other words, these panels provide information as to whether the effect of a given variable is higher or smaller than expected given taxonomic composition. **c**) Variation in the magnitude of the standard effect size (SES) value along the observed gradient. In other words, these panels provide information as to whether the effect of a given variable in determining trait dispersion changes along the gradient. In **b** and **c**, significant effects (Rank <0.025 |> 0.975) are highlighted with a darker purple.

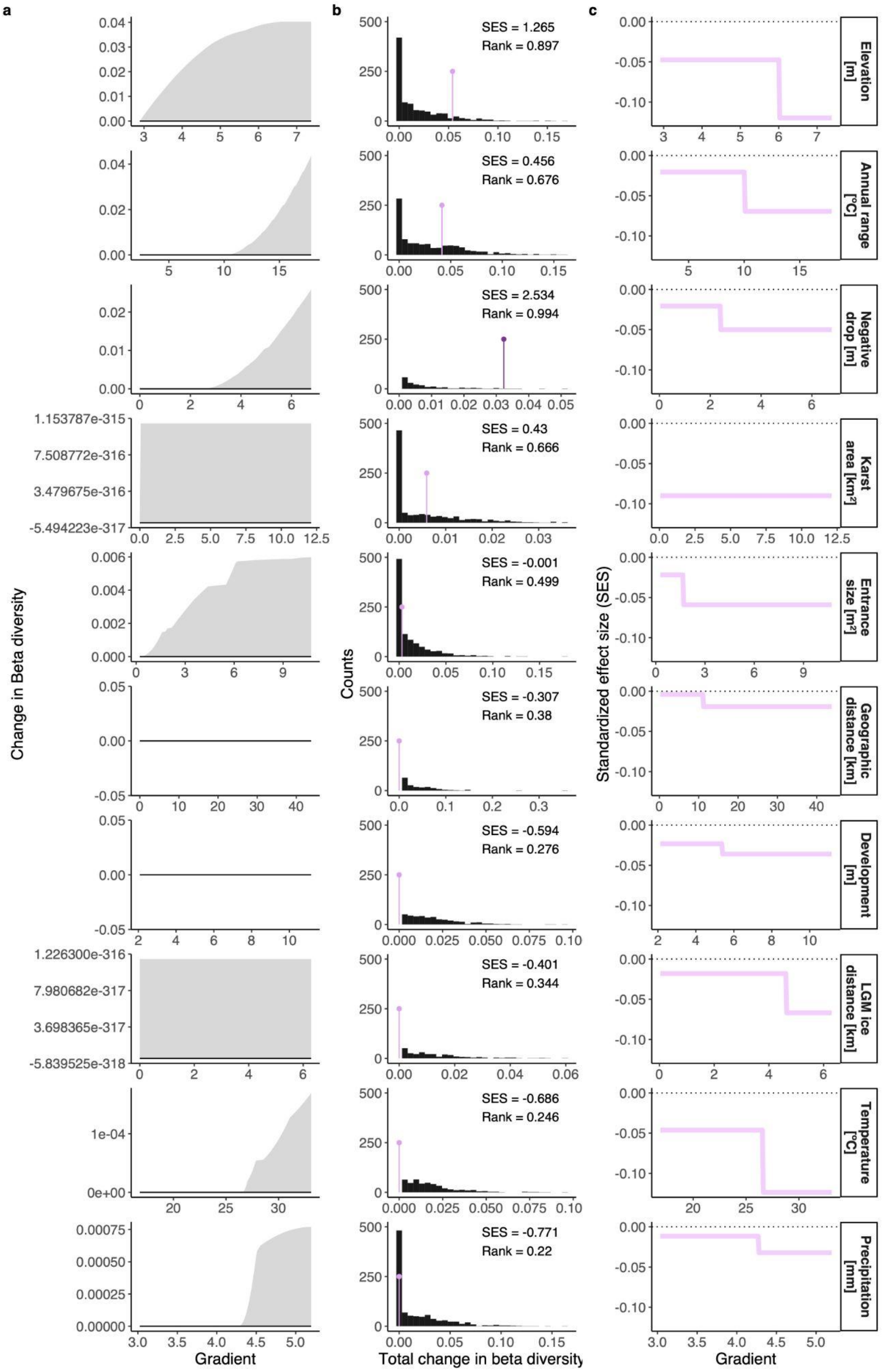


**Figure S2.** Results of Bayesian bootstrap generalized dissimilarity modeling for change in the richness component of functional β-diversity ($\beta_{richness}$) of subterranean spider communities across Europe. Variables are sorted by their contribution (on top: highest contribution). **a**) Fitted I-splines (partial regression fits) for the considered environmental and geographic gradients. The maximum height reached by each curve indicates the total amount of compositional turnover explained by that variable (holding all other variables constant), whereas the shape of each spline indicates how the rate of compositional turnover varies along the gradient. **b**) Distribution of expected values (histogram) versus the observed value (colored line) of each environmental and geographic gradient, based on null modeling (999 iterations). In other words, these panels provide information as to whether the effect of a given variable is higher or smaller than expected given taxonomic composition. **c**) Variation in the magnitude of the standard effect size (SES) value along the observed gradient. In other words, these panels provide information as to whether the effect of a given variable in determining trait dispersion changes along the gradient. In **b** and **c**, significant effects (Rank <0.025 |> 0.975) are highlighted with a darker purple.
